# Passive inertial damping improves high-speed gaze stabilization in hoverflies

**DOI:** 10.1101/2021.12.11.472180

**Authors:** Ben J Hardcastle, Karin Bierig, Francisco JH Heras, Kit D Longden, Daniel A Schwyn, Holger G Krapp

## Abstract

Gaze stabilization reflexes reduce motion blur and simplify the processing of visual information by keeping the eyes level. These reflexes typically depend on estimates of the rotational motion of the body, head, and eyes, acquired by visual or mechanosensory systems. During rapid movements, there can be insufficient time for sensory feedback systems to estimate rotational motion, requiring additional mechanisms. Solutions to this common problem are likely to be adapted to an animal’s behavioral repertoire. Here, we examine gaze stabilization in three families of dipteran flies, each with distinctly different flight behaviors. Through frequency response analysis based on tethered-flight experiments, we demonstrate that fast roll oscillations of the body lead to a stable gaze in hoverflies, whereas the reflex breaks down at the same speeds in blowflies and horseflies. Surprisingly, the high-speed gaze stabilization of hoverflies does not require sensory input from the halteres, their low-latency balance organs. Instead, we show how the behavior is explained by a hybrid control system that combines a sensory-driven, active stabilization component mediated by neck muscles, and a passive component which exploits physical properties of the animal’s anatomy—the mass and inertia of its head. This adaptation requires hoverflies to have specializations of the head-neck joint that can be employed during flight. Our comparative study highlights how species-specific control strategies have evolved to support different visually-guided flight behaviors.

**SIGNIFICANCE STATEMENT:** Across the animal kingdom, reflexes are found which stabilize the eyes to reduce the impact of motion blur on vision—analogous to the image stabilization functions found in modern cameras. These reflexes can be complex, often combining predictions about planned movements with information from multiple sensory systems which continually measure self-motion and provide feedback. The processing of this information in the nervous system incurs time delays which impose limits on performance when fast stabilization is required. Hoverflies overcome the limitations of sensory-driven stabilization reflexes by exploiting the passive stability provided by the head during roll perturbations with particularly high rotational kinematics. Integrating passive and active mechanisms thus extends the useful range of vision and likely facilitates distinctive aspects of hoverfly flight.

## INTRODUCTION

Agile flight maneuvers require a keen sense of vision, but without compensatory mechanisms visual processing would be severely impaired during fast movement ^1^. Gaze stabilizing reflexes have evolved in many animals, which reduce motion blur and keep the eyes and visual coordinates aligned with the horizon ^2–4^. When the eyes are fixed to the head or have a limited range of motion—as in barn owls and many flying insects—head movements play a pivotal role in stabilizing gaze. The actuation of compensatory head movements is a sophisticated calculation which must handle the different time delays of the various sensory feedback systems involved, as well as taking into account the mechanical properties of the head and the range of movements the neck muscles can actuate ^5^.

Sensory feedback systems with low latency are particularly valuable for stabilizing gaze during high-speed maneuvers, and in flies (Diptera) the halteres fulfill this role ^6^. The halteres are a pair of club-shaped appendages on the thorax which have evolved from a rear pair of wings and act as the principal balance organs, sensing the angular velocity of the body ^7–11^. In addition, the angular position of the head relative to the body is monitored by proprioceptors, and the motion of the head is measured visually through slower processing dependent on the compound eyes^12,13^. Many fly species also have ocelli, a set of three small, simple lens eyes on the top of the head which rapidly detect changes in orientation through differential illumination ^14,15^.

Since dipterans are diverse and exhibit different styles of flight and specializations of their sensory systems ^16–20^, we hypothesized that gaze stabilization would also demonstrate species-specific adaptations, whose mechanisms would reveal solutions to motor control tasks at the limits of temporal precision. To test our hypothesis, we compared species from three families with contrasting behaviors: blowflies (Calliphoridae), horseflies (Tabanidae), and hoverflies (Syrphidae) ^16^.

Blowflies form the basis of our comparison, since the gaze stabilization system which compensates for body-roll has been extensively studied in these species ^12,13^. Their flight is characterized by high-acceleration body saccades and banked turns ^21^, as well as high-speed aerial pursuits launched from a perch ^22–24^ and low-speed circling around food sources prior to landing.

Female horseflies, on the other hand, use polarized light cues to detect hosts from a distance across open fields and exhibit direct flights toward them at speed ^25–27^. Although typically larger than blowflies, these insects are capable of agile aerial maneuvers ^28^ and males are often observed hovering in swarms for the purposes of mating ^29–31^. The horsefly species we investigate here, *Tabanus bromius*, lack functional ocelli— the simple eyes found dorsally on the head of blowflies and hoverflies.

Hoverflies, while bearing similarities in many flight maneuvers to blowflies, also hover with exquisite control for extended periods, and are notable for darting and shadowing conspecifics, as well as their ability to fly backwards while hovering ^19,22,32,33^. We initially compared roll gaze stabilization performance across the three dipteran families and searched for differences which might reflect their flight behavior.

## RESULTS

### Hoverfly gaze stabilization improves at high speeds

To evaluate gaze stabilization performance across species, we used a tethered-flight paradigm and induced oscillations of the thorax around the longitudinal (roll) axis of the animal (Fig. 1A left). Experiments were captured on a high-speed camera, and the absolute roll angles of the head and thorax were measured relative to the vertical axis in each frame (Fig. 1A right). We applied a ±30°sinusoidal chirp stimulus which varied the oscillation frequency of the thorax over time: first increasing linearly from 0 to 20 Hz in 5 s, then decreasing again from 20 to 0 Hz in 5 s. Perfect gaze stabilization would result in rotations of the head equal and opposite to those of the thorax, with zero delay, and would be reflected by a motionless head in the camera view.

**Figure 1.**
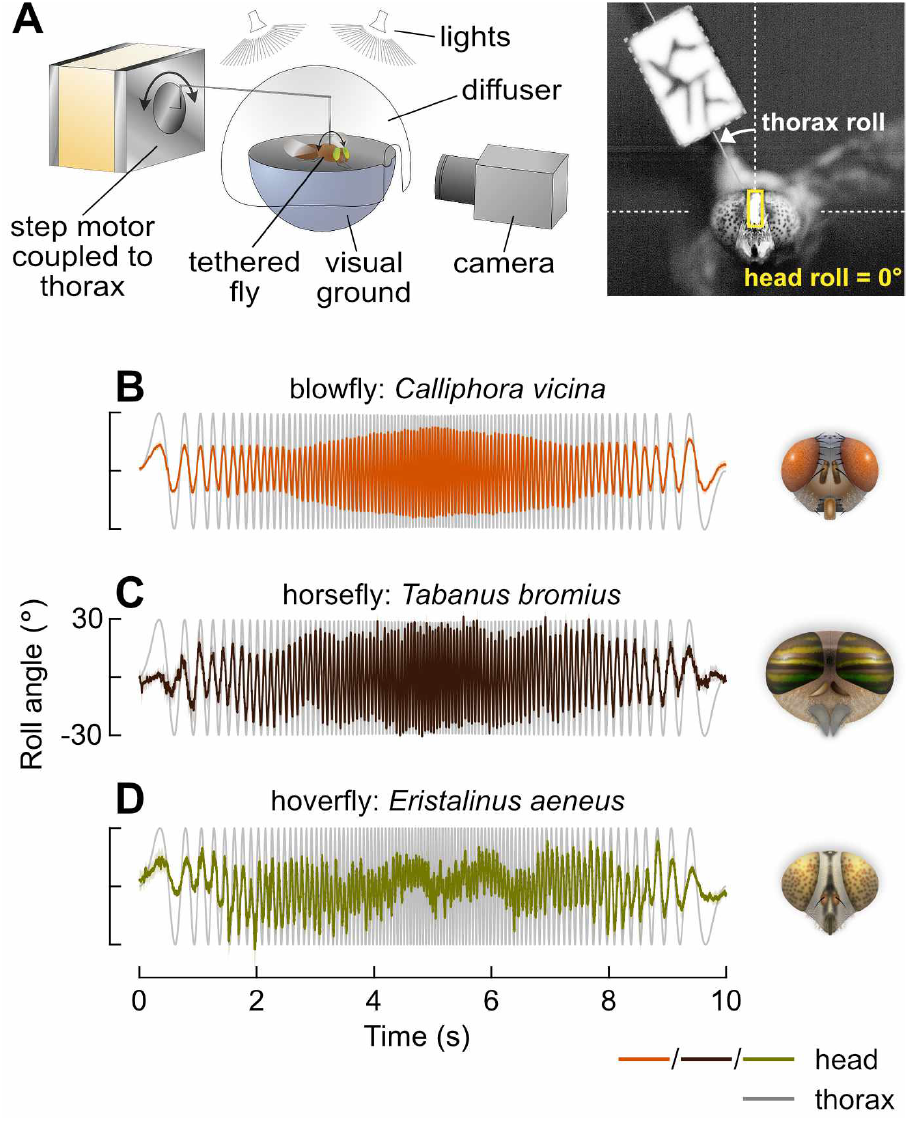
Hoverfly gaze stabilization performance improves at high speeds. **A:** Experimental setup (left). Flies were tethered at the thorax to a step motor via a piece of cardboard. Oscillations of the motor simulated thorax roll perturbations of the fly. Diffuse light was delivered from the dorsal hemisphere while a dark ground in the ventral hemisphere provided a horizon as a visual reference for stabilization. A high-speed video camera captured the resulting compensatory rotations of the head (right). Painted markers on the head and tether aided tracking. **B:** Average time-series from experiments using a sinusoidal chirp stimulus, for the blowfly (*C. vicina*). The stimulus oscillated the thorax (gray trace) with a time-varying frequency profile. The absolute angle of the fly’s head (color trace) is overlaid, demonstrating a stabilization effort which generally reduced the roll amplitude of the head in all species. Perfect stabilization would appear as a flat line at 0° and no stabilization effort would result in the head angle following the thorax angle, oscillating at ±30°. Traces show mean head roll angle across flies. Shaded area shows mean ± standard error (8 flies). **C:** As in **A**, for the horsefly (*T. bromius*: 4 flies). **D:** As in **A**, for the hoverfly (*E. aeneus*: 6 flies).

At low frequencies, the gaze stabilization reflex in each species is effective at reducing the motion of the head compared to the motion of the thorax (0–1 s, Fig. 1B–D). But as frequency increases, stabilization performance decreases: the rotational speeds exceed the operating range of the sensory systems contributing to the stabilization reflex and the amplitude of head roll motion becomes progressively larger. For the blowfly, *Calliphora vicina*, head roll amplitude continued to grow until the thorax oscillations slowed down at the mid-point of the experiment (5 s, Fig. 1B, Movie 1). The same occurred for the horsefly, *Tabanus bromius*, where head roll approached the ±30° motion of the thorax, indicating an almost completely ineffectual stabilization reflex (Fig. 1C, Movie 2) (note that ‘amplitude’ refers here to the motion of the head as measured from the camera frame of reference: as stabilization performance decreases, the compensatory movements of the head relative to the thorax become smaller, resulting in increasing amplitude in the camera frame).

This negative relationship between frequency and gaze stabilization performance, above a certain frequency optimum, has previously been observed in flies ^34,35^, as well as in other animals (flying insects ^36,37^, birds ^38^, fish ^39^, reptiles and amphibians ^40^, crustaceans ^41^, and mammals ^42,43^—including humans ^44^). Although it appears to be a common property across taxa—a consequence of the limited operating range of an animal’s visual and mechanosensory systems—the gaze stabilization performance of the hoverfly, *Eristalinus aeneus*, showed a different dependence on frequency. At the highest frequencies, the hoverfly’s head roll amplitude is smaller than at intermediate frequencies (Fig. 1D, Movie 3). It is also much reduced compared to the blowfly and horsefly.

To confirm that this effect was not caused by the time-varying frequency sweep contained within the chirp stimulus, we performed similar experiments using constant-frequency stimuli. Again, we observed that head roll amplitude grew larger with frequency for the blowfly and horsefly (Fig. 2A,B). For the hoverfly, head roll amplitude grew from an average of ±8° at 1 Hz to ±18° at 10 Hz—a similar increase to the other species (Fig. 2C). However, as in the chirp experiment, head roll amplitude then became smaller again at the highest speeds tested, falling to around ±10° at 20 Hz.

**Figure 2.**
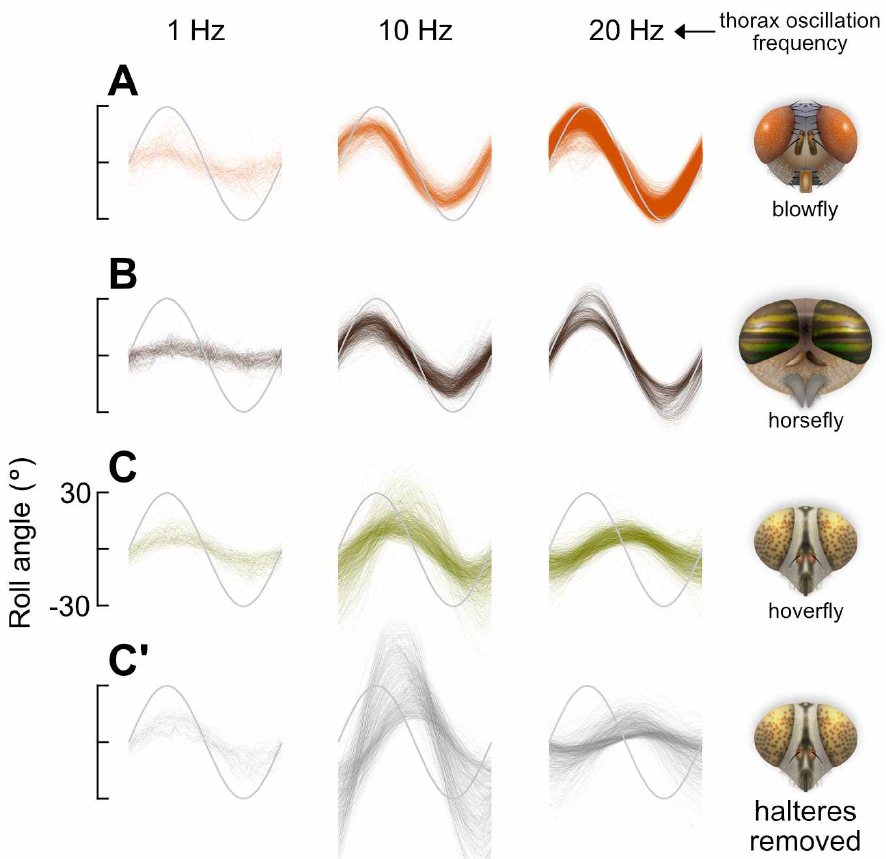
High-speed stabilization in hoverflies does not require haltere input. **A:** Time-series of head roll angle (color traces) in response to individual cycles of constant-frequency sinusoidal oscillations of the thorax (gray traces), for the blowfly (*C. vicina*: 5–13 flies). Perfect stabilization would appear as a flat line at 0° and no stabilization effort would result in the head angle following the thorax angle, oscillating at ±30°. **B:** As in **A**, for the horsefly (*T. bromius*: 8 flies). **C:** As in **A**, for the hoverfly (*E. aeneus*: 6 flies). **C’:** Responses of the animals shown in **C** after removing the halteres. At 10 Hz (center), the motion of the head increased compared to the intact response, while at 1 Hz (left) and 20 Hz (right), the motion of the head was comparatively unaffected.

### High-speed stabilization in hoverflies does not require haltere input

At high speeds, the predominant sensory input to gaze stabilization in the blowfly is provided by the halteres ^6^. Are hoverfly halteres simply tuned to detect higher frequency oscillations than those of the blowfly and horsefly? When we repeated the previous experiment in the hoverfly *E. aeneus* after removing the halteres, head roll motion at the intermediate 10 Hz frequency was increased greatly compared to the intact response (Fig. 2C,C’ center). Indeed, head roll oscillations became larger than those of the thorax, consistent with a framework in which sensory input from the halteres is crucial for effective gaze stabilization ^35^. Contrary to this notion, however, increasing the frequency to 20 Hz with the halteres removed elicited a more effective stabilization of the head: compared to the intact condition, haltere removal had no discernible effect on either the amplitude or the phase of head roll motion at 20 Hz (Fig. 2C,C’ right).

Frequency response plots for each animal illustrate the differences in their gaze stabilization behavior (Fig. 3A–C). Linear gain—a proxy for performance—falls to around zero at 25 Hz in the response of the intact blowfly, and at 10 Hz with its halteres removed (Fig. 3A). For the horsefly, zero gain occurs at approximately the same frequencies as for the blowfly (Fig. 3B). A large negative gain is also observed at >15 Hz, which may be interpreted as head roll motion being increased by the gaze stabilization system at high speeds, rather than reduced: as the period of the stimulus becomes shorter, the relatively constant delay in visual feedback grows as a proportion of each stimulus cycle duration (phase lag), ultimately causing compensatory rotations of the head to be actuated at a phase which adds to the thorax roll instead of reducing it.

**Figure 3.**
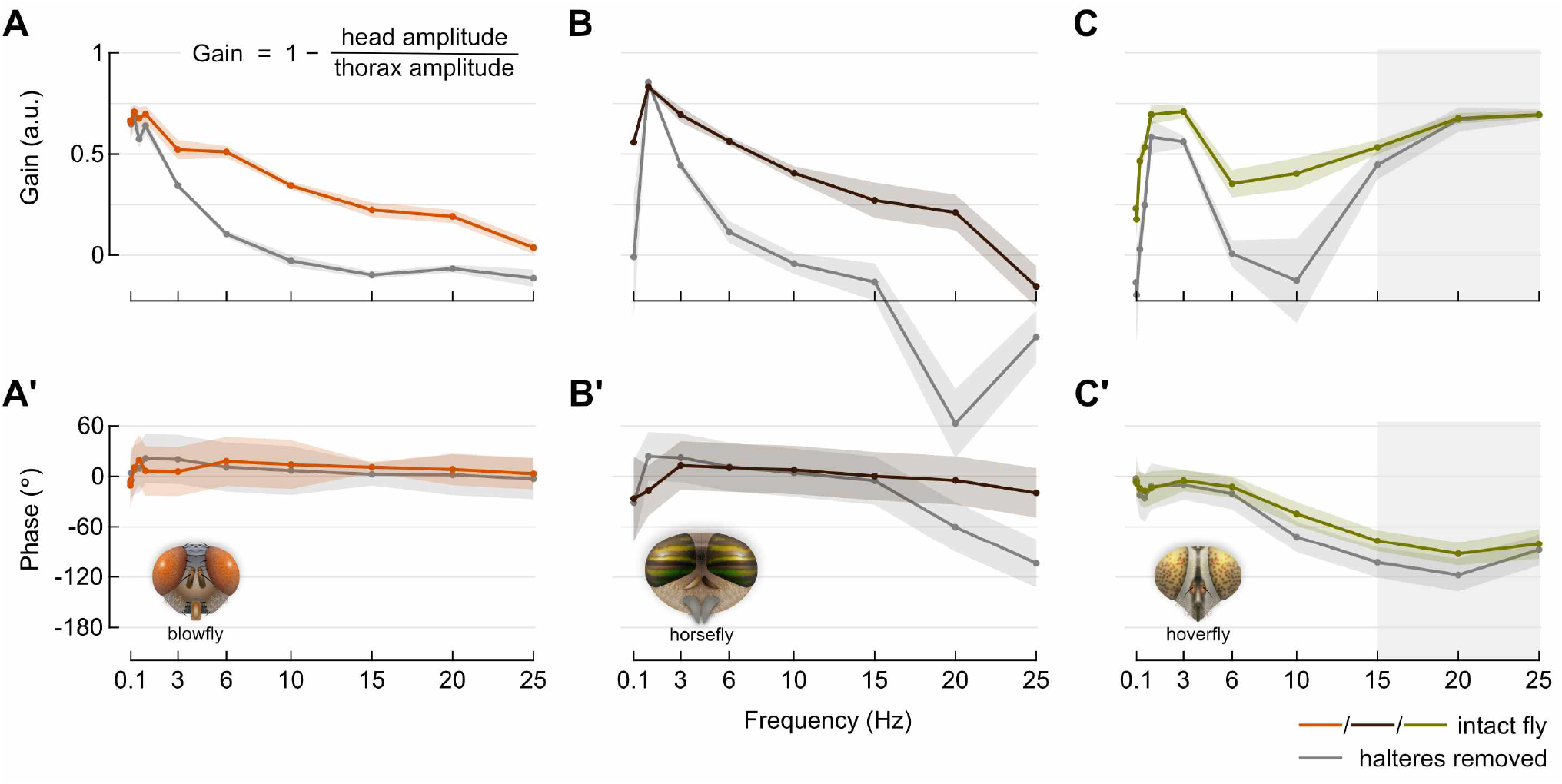
Gain and phase of head roll frequency-response. **A:** Average gain of the head roll response for the intact blowfly (color trace) and after removing the halteres (gray trace). Data obtained from experiments using constant-frequency sinusoidal stimuli. Shaded area shows mean ± standard error (*C. vicina*: 5–13 flies). Head and thorax amplitudes are measured from the camera frame of reference, as in Fig. 1. **A’:** Corresponding phase angle of head roll response for the data shown in **A**. **B:** As in **A**, for the horsefly (*T. bromius*: 8 flies). Negative gain values at 20 and 25 Hz with the halteres removed indicate increased motion of the head relative to the motion of the thorax. **C:** As in **A**, for the hoverfly (*E. aeneus*: 6 flies). Gray shaded area indicates high-frequency range in which gain is unaffected by removing the halteres (gain: *P* = 0.33 at 15 Hz, *P* = 0.53 at 20 Hz, *P* = 0.33 at 25 Hz, Wilcoxon rank-sum test). **C’:** Gray shaded area indicates high-frequency range in which gain is unaffected by removing the halteres (phase: *P* < 0.005 at 15 Hz, *P* = 0.041 at 20 Hz, *P* = 0.47 at 25 Hz, Wilcoxon rank-sum test).

For the hoverfly, gain does not fall to zero with the halteres intact (Fig. 3C): the negative trend with frequency is clearly reversed between 3 Hz and 6 Hz. With its halteres removed, only frequencies <15 Hz are impacted: at 15, 20 and 25 Hz, we found no significant difference in gain versus the intact condition (Fig. 3C gray shaded area). At 0.3 Hz and below, we noted that the low speed oscillations often did not elicit a large stabilization effort in the hoverfly, resulting in gains well below 0.5 in both conditions.

Two different gaze stabilization behaviors are thus evident in the hoverfly frequency response: a lower-speed regime which requires mechanosensory input from the halteres and a higher-speed regime which operates independently of the halteres. Is it possible that other sensory inputs are contributing to this higher-speed regime? If the head is sufficiently stabilized, the speeds of visual motion may be within the operating range of the compound eyes—one of the key benefits of a stabilization reflex—which would allow them to contribute to the reflex itself, as they likely do at lower speeds (see gain at 1 Hz and 3 Hz with halteres removed, Fig. 3C). However, the motion applied to the thorax at 25 Hz exceeds 5000°s^−1^, and it is implausible that the visual system alone is responsible for the stabilization observed.

The phase lag (delay) calculated for the hoverfly head response was considerably longer than for the other flies (Fig. 3A’– C’, Fig. 2C). Combined with a gain close to unity, a long phase lag could cause the stabilization system to increase the motion of the head, rather than reduce it. We therefore asked how effective hoverfly gaze stabilization is at reducing head motion to speeds which are within the operating range of the compound eyes.

### Gaze stabilization is effective over a wider dynamic range in hoverflies than in other flies

At each frequency tested, we found the probability distribution of retinal slip-speeds experienced by each fly, i.e. the speed of visual motion across the eyes (Fig. 4A–C). For each distribution we also marked the maximum slip-speed that would typically be experienced if no stabilization effort were made (slip-speed = thorax speed).

**Figure 4.**
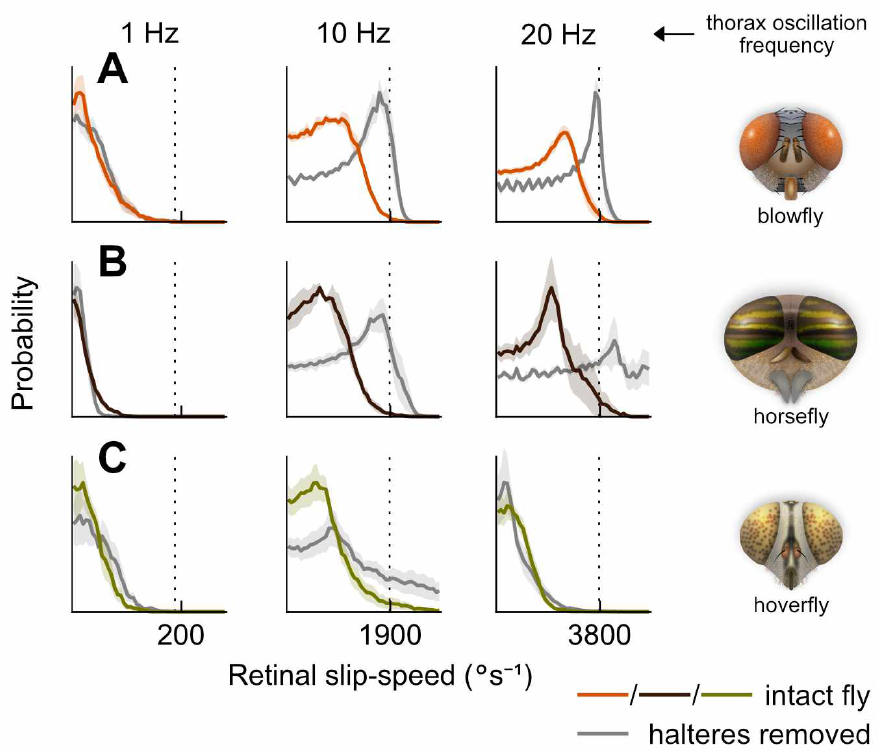
Slip-speed distributions demonstrate effectiveness of stabilization at different frequencies. **A:** Normalized probability distribution of visual slip experienced by the intact blowfly (color traces) during constant-frequency sinusoidal oscillations, and for the same animals after removing the halteres (gray traces). Shaded area shows mean ± standard error (*C. vicina*: 5–13 flies). Vertical dashed line indicates theoretical maximum slip-speed experienced with no stabilization effort (i.e. head angle = thorax angle). **B:** As in **A**, for the horsefly (*T. bromius*: 8 flies). **C:** As in **A**, for the hoverfly (*E. aeneus*: 6 flies).

As expected for the blowfly and horsefly, the peak (mode) of each distribution shifts progressively further towards higher slip-speeds with increasing stimulus frequency, and upon removal of the halteres (Fig. 4A,B, Fig. 5A,B). Based on typical measurements of the compound eye geometry and photoreceptor response characteristics in blowflies and hoverflies, we estimated the slip-speed at which motion blur would begin to degrade spatial information to be between 100–200°s^−1^ (see Materials and Methods). The blowfly and horsefly both pass this limit, and are far beyond it at 15 Hz, or 10 Hz with their halteres removed (Fig. 5A,B), while slip-speed in the hoverfly plateaus just above this approximate limit for the intact animal (Fig. 5C). With the halteres removed, the mode of the slip-speed distribution exceeds 1000°s^−1^ in the hoverfly at 10 Hz, but is brought under the limit at higher frequencies. We conclude that gaze stabilization in *E. aeneus* is effective across a wider dynamic range than in the other two species, and likely reduces head motion to be within, or close to, a range in which visual information is only mildly affected by motion blur.

**Figure 5.**
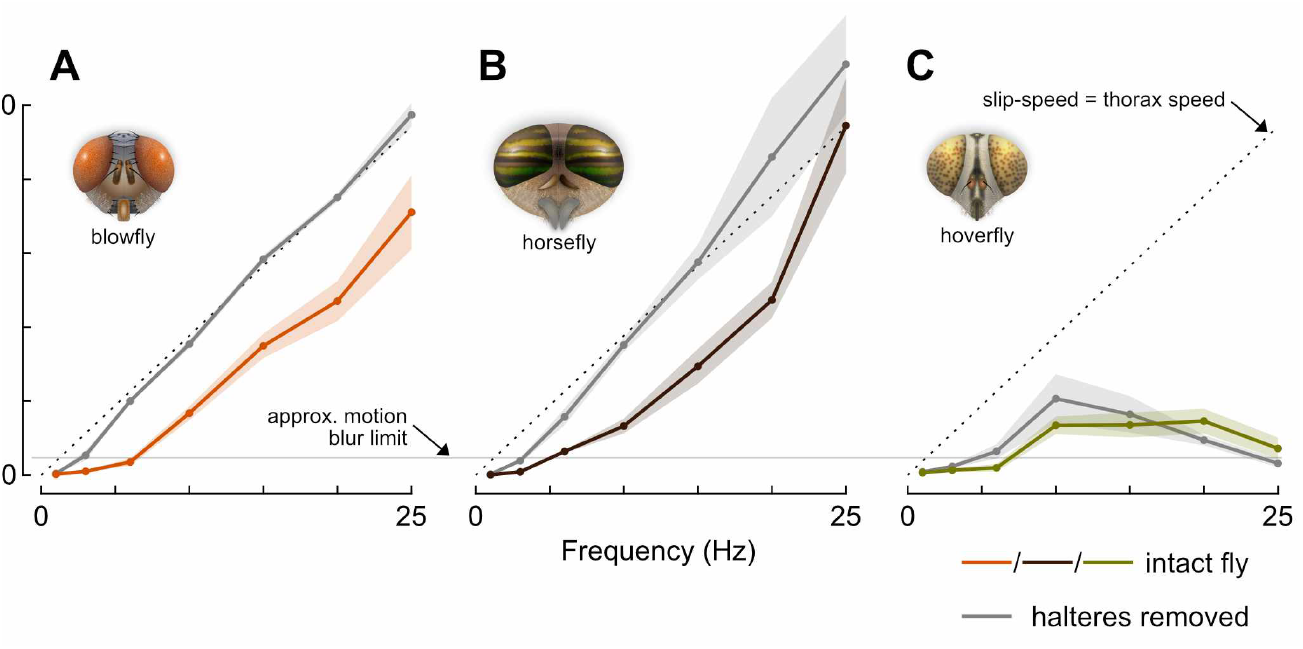
Gaze stabilization is effective over a wider dynamic range in hoverflies than in other flies. **A:** Mode (peak) values of the probability distributions of visual slip experienced by the intact blowfly (color trace) during constant-frequency sinusoidal oscillations, and for the same animals after removing the halteres (gray traces). Shaded area shows mean ± standard error (*C. vicina*: 5–13 flies). **B:** As in **A**, for the horsefly (*T. bromius*: 8 flies). **C:** As in **A**, for the hoverfly (*E. aeneus*: 6 flies).

### Hoverfly head-neck joint facilitates stabilization through inertial damping

We next asked whether the high-speed gaze stabilization behavior is unique to *E. aeneus*, and how it might function. To answer these questions we turned to two other members of the Syrphidae family: the common drone fly, *Eristalis tenax*, and the marmalade fly, *Episyrphus balteatus*. In both of these hoverfly species, we found stabilization behavior in response to the chirp stimulus which was qualitatively similar to *Eristalinus aeneus*, with a reduction in head roll amplitude at high frequencies (Fig. 6A,B). This finding suggests that a similar mechanism may facilitate high-speed stabilization across hoverflies.

**Figure 6.**
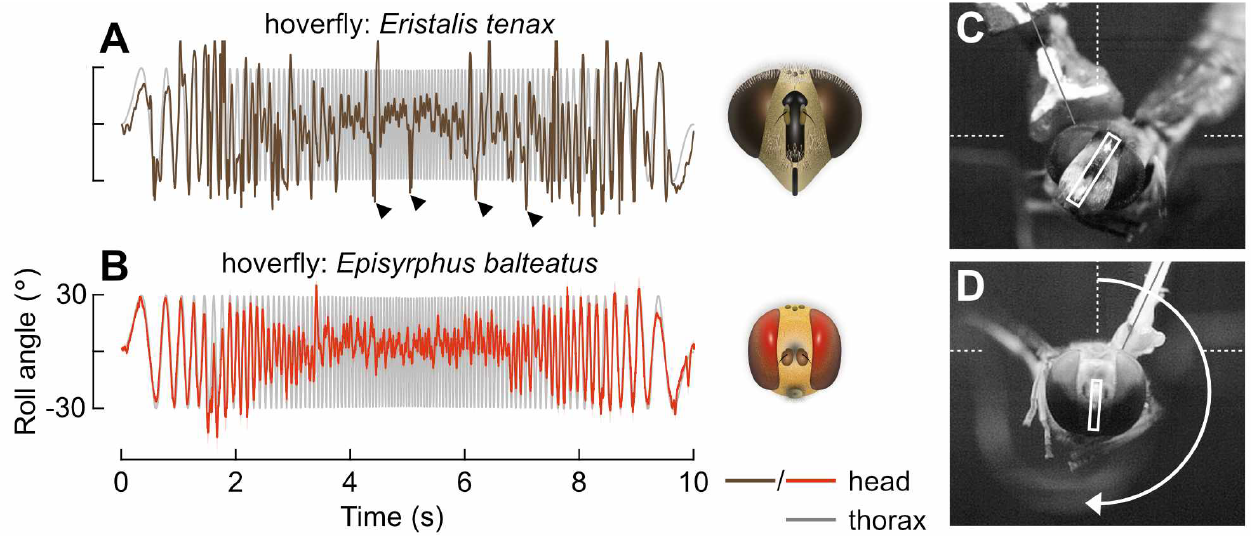
Specializations of the head-neck motor system in hoverflies may enable inertial stabilization. **A:** Time-series from a single chirp experiment for a second species of hoverfly (*E. tenax* : 1 fly). Arrowheads indicate large angle, spontaneous roll rotations of the head which were uncorrelated with the stimulus. **B:** Average time-series from chirp experiments for a third species of hoverfly. Trace shows mean head roll angle across flies. Shaded area shows mean ± standard error (*E. balteatus*: 13 flies). **C:** Frame capture of *E. tenax* chirp experiment during brief stabilization of the head at an offset roll angle (see Movie 4). **D:** Frame capture of *E. balteatus* experiment showing inversion of the head (see Movie 5).

During experiments with syrphids, we observed a number of intriguing features of head movements that were not present in the calliphorid and tabanid species we investigated—behaviors which indicated specializations of the hoverfly neck motor system. First, we observed an apparent loosening, or relaxation, of the head-neck joint, which resulted in a distinctive ‘wobble’ of the head at intermediate to high frequencies (10–20 Hz). Head wobble events were visible in all three hoverfly species as small amplitude motion of the head (less than a few degrees) at frequencies far higher than the thorax oscillation. A distinguishing feature of head wobble was periodic motion, usually around the pitch or yaw axes, with a noticeable settling time (Movie 3–Movie 5). These events typically occurred upon reversal of thorax motion. In each species, the head wobble gave the impression of a mass rotating on a loose pivot, i.e. the head-neck joint exhibited lower stiffness, damping and friction than the blowfly and horsefly species, which lacked such wobble (Movie 1, Movie 2). Small mechanical juddering induced by the step-motor at the extreme of each cycle appeared to shake the animals, and in hoverflies the head wobbled as a result.

Next, we observed occasional periods of static roll angle offset, during which the hoverfly’s head was stabilized and relatively free of motion, but not in the default upright orientation. Rather, the head remained rolled at an offset angle (approximately 30–60°) for one or more cycles of the stimulus (Fig. 6C, Movie 3–Movie 5). Erroneous sensory information could explain this observation: the prosternal organs, for example, detect head angle relative to the thorax and affect static roll offsets in blowflies ^45^. However, the kinematics of the head were qualitatively different to those at low frequencies (<10 Hz) or in the blowfly or horsefly, and gave the impression that head movements were not under active control of the neck muscles during periods of offset (Movie 3).

Finally, in *E. tenax* and *E. balteatus*, large roll rotations of the head occurred during experiments (Fig. 6A arrowheads). In *E. balteatus*, these rotations were often extreme, completely inverting the head (Fig. 6D, Movie 5). The rotations occurred spontaneously, in that they were seemingly uncorrelated with the motion of the thorax. Notably, the head appeared to rotate until reaching a mechanical limit with sufficient force that it rebounded, again indicating low damping in the head-neck joint. In addition, the head did not rapidly return to an upright orientation upon rebound, as would be expected if the head-neck joint exerted an elastic restoring force, but returned slowly, wobbled, or remained at an offset, suggesting low torsional stiffness (Movie 5).

Based on these observations, we propose that active control of the neck muscle system may at times be selectively disabled, allowing mechanical forces acting on the head to passively influence its motion. In this state, it is possible that the inertia of the head could damp forced rotations of the thorax and stabilize the default orientation of the head without sensory input.

### A head-neck model captures high-speed hoverfly stabilization behavior

Could inertial damping explain the stabilization behavior observed in hoverflies? Modeling a purely passive, frictionless head-neck joint system with reduced torsional stiffness and damping constants shows that head roll amplitude does indeed decrease with frequency in response to a chirp stimulus (Fig. 7A,B), strongly resembling the behavioral response observed in hoverflies at high frequencies (Fig. 1D, Fig. 6A,B).

**Figure 7.**
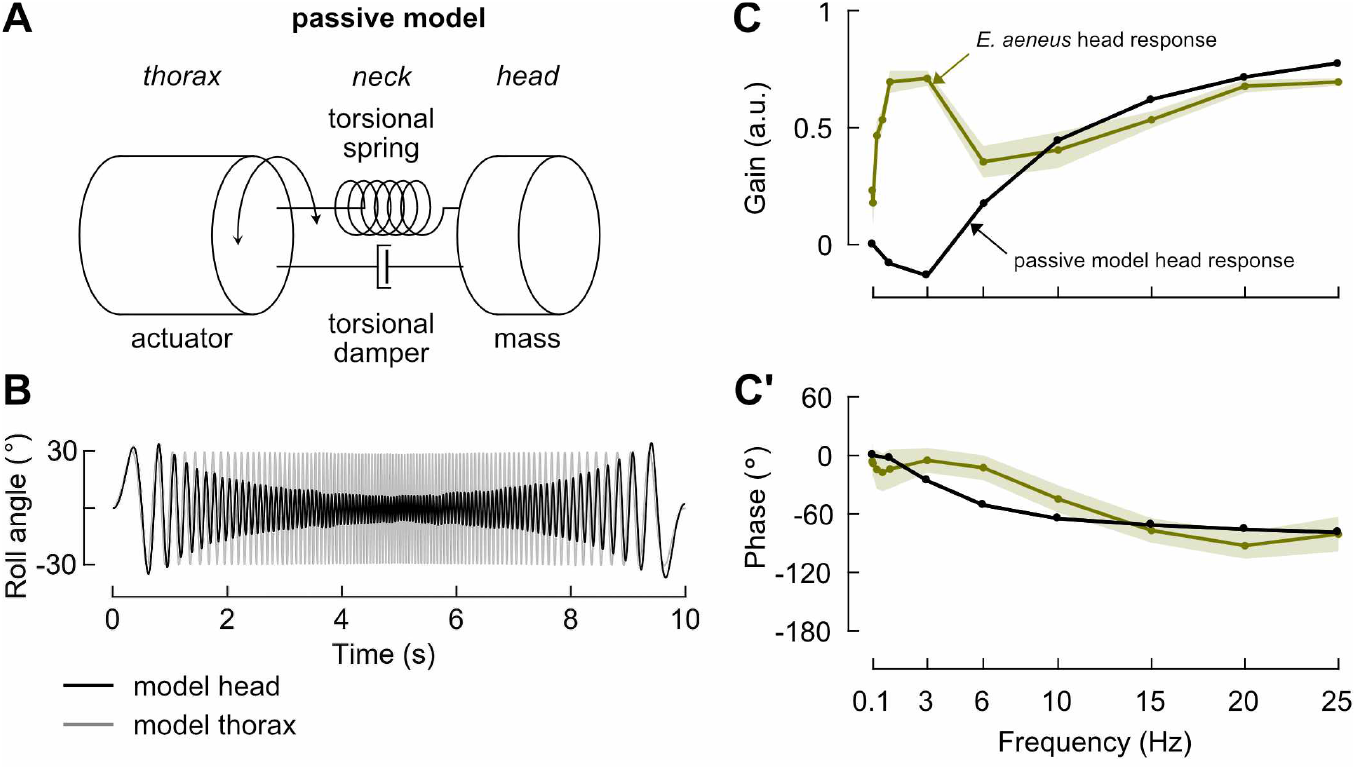
A head-neck model with low torsional stiffness captures high-speed hoverfly stabilization behavior. **A:** Diagram of a passive mechanical model of the hoverfly head, neck and thorax. The neck is modeled by a torsional spring and damper, and couples the mass of the head to the thorax, which is driven by forced oscillations. **B:** Time-series from a simulated experiment using a sinusoidal chirp stimulus applied to the passive model shown in **A**. The stimulus oscillated the model thorax (gray trace) with a time-varying frequency profile. The absolute angle of the model head (black trace) is overlaid, demonstrating a completely passive, inertial stabilization which reduced the roll motion of the head relative to the thorax. Perfect stabilization would appear as a flat line at 0°. **C:** Average gain of the head roll response for the model head (black trace). Data obtained from simulated experiments using constant-frequency sinusoidal stimuli. The intact hoverfly data (yellow trace) are replot from Fig. 3C for comparison. **C’:** Corresponding phase angle of head roll response for the data shown in **C**.

Simulations of constant-frequency oscillations further demonstrate that at low frequencies—up to around 3 Hz—the forces on the head are insufficient for inertial damping to stabilize it, and the motion of the head approximately follows the thorax, which results in gains close to zero (Fig. 7C, Fig. S1A). For the hoverfly *E. aeneus*, gains are higher than predicted by the passive model in the range 0.06–3 Hz (Fig. 7C), indicating an active gaze stabilization reflex that depends on sensory input. Where the gain of the hoverfly response drops between 3–10 Hz, the gain in the passive model increases as inertia begins to affect head motion. Between 10–25 Hz, the gain and phase of the passive modeled response closely match the hoverfly data (Fig. 7C, C’), with a similar plateau in slip-speeds at around 600°s^−1^ (Fig. S1C, Fig. 5C). This leads us to conclude that passive, inertial damping alone, with no sensory input, could provide effective gaze stabilization at high speeds, provided that the stiffness and damping of the head-neck joint are appropriately low.

## DISCUSSION

Here we have presented lines of evidence which support a view of gaze stabilization through inertial damping in hoverflies. This passive behavior enables effective stabilization of the head and eyes while the thorax is free to roll at extremely high angular velocities and accelerations. While we uncovered this behavior in a tethered-flight paradigm with a motor actuating roll oscillations of the thorax, we expect that it would be similarly activated in response to external disturbances in free-flight, such as wind gusts.

The repetitive, oscillatory motion of the sinusoidal stimuli used in our experiments is clearly different to that of a wind gust, and investigating responses to an abrupt, step-like rotation of the thorax would have been desirable in this sense. The prohibitively high inertia of the motor used in our setup did not allow us to generate roll accelerations well approximating a step function. Goulard et al. ^46^, however, were able to induce step-like thorax rolls in *E. balteatus*. In their study, the hoverfly head showed an amount of overshoot upon step rotations which is congruous with the low stiffness and damping of the neck which we propose allows inertia to stabilize the head.

### Inertial gaze stabilization in the context of hoverfly flight behavior

Inertial gaze stabilization, which was unaffected by removing the mechanosensory input from the halteres, was observed in our experiments at oscillation frequencies of 15 Hz and greater. At 15 Hz, the maximum angular velocity applied to the thorax was around 2800°s^−1^, and maximum acceleration was around 2 × 10^5^°s^−2^. Do hoverflies actually encounter roll rotations with comparable kinematics during flight? Previous studies which have captured the free-flight behavior of hoverflies (*E. tenax, E. balteatus*, and various other species) either did not resolve or report roll rotations of the thorax ^22,47–51^, but similar experiments with blowflies (*C. vicina*) recorded roll velocities in excess of 2000°s^−1^ and accelerations on the order of 10^5^°s^−2^ during fast U-turn maneuvers and saccades^21^. Meanwhile, landing maneuvers made by *C. vomitoria* can involve a rapid inversion of the body about the roll axis, with velocities approaching 6000°s^−1 52^. These volitional maneuvers took place in relatively small, confined arenas, and even higher values may well be expected in the wild.

However, our experiments captured reflexive behavior in response to roll rotations caused by an external disturbance, rather than voluntary movements. One study examining the impact of such external perturbations on insect flight demonstrated that hovering bees (*Apis melifera*) are capable of rapid recovery from a wind gust which caused roll rotations with similar kinematics ^53^. In another study, a sudden free-fall situation was imposed on stationary hoverflies (*E. balteatus*) hanging from a ceiling, which induced a righting maneuver to recover from the tumble ^54^. In these experiments, extremely high roll rates of over 10 × 10^3^°s^−1^ were recorded. The animals’ ability to regain stability after such perturbations makes it reasonable to assume that they regularly encounter such excessive attitude changes during natural flight in turbulent conditions.

Why, then, does it appear that hoverflies employ inertial gaze stabilization while other highly maneuverable flies like blowflies do not? We find clues to answer this when we consider the distinguishing flight behavior of hoverflies—namely, hovering, for the purpose of visiting flowers, guarding territory and seeking mates. While hovering, flies may be particularly susceptible to being rolled by gusts of wind. Lateral instability is higher when hovering than during forward flight ^55,56^ and angular velocities around the roll axis are typically higher than those around pitch or yaw for an insect flying in turbulent conditions, due to a smaller moment of inertia ^57^. Hoverflies also seem to be equipped for more agile flight than the other dipteran families we investigated here: wide-field motion sensitive visual neurons in hoverflies respond more rapidly than the homologous neurons in *Calliphora* spp., for example ^19^, and are greater in number in each individual animal ^18,58^. They also maintain sensitivity across a wider range of temporal frequencies of image motion ^17^.

Hovering in hoverflies may therefore be particularly demanding in terms of flight maneuvers and stabilization reflexes. The gaze stabilization system in other flies might not be required to operate at a dynamic input range that includes such high angular accelerations that may occur while holding a hovering position for extended periods or during the initial phase of an aerial pursuit. Another possibility is that the visually-guided behaviors which hovering flight supports are also highly demanding in hoverflies and necessitate this alternative stabilization method. For example, the detection of conspecifics before initiating aerial pursuits from hovering likely requires near-constant high-acuity, stabilized vision, which may be a less demanding sensorimotor task for ground-launched pursuits. Likewise, the flight reflexes to recover from a gust-induced tumble may tolerate some degree of brief motion blur due to passive stability afforded by the body and wings.

### Anatomical specializations of the head-neck joint

How could the head-neck joint work in hoverflies to enable inertial stabilization? First, we posit that a flexible joint is required, with lower stiffness and damping than the equivalent joint in the species of blowfly or horsefly investigated here. Low friction in the joint is also necessary, to allow the head to effectively spin freely while the thorax rotates. When allowed to spin freely, rotations of the thorax are decoupled from the head. The head then tends to remain in a default orientation as a result of its inertia—at least, for a certain range of rotational accelerations.

Below this range, the effect of inertia is insufficient to overcome the torsional stiffness of the joint. The head is then more strongly influenced by rotations of the thorax and inertia provides little stabilization, as seen in the response of a purely passive model of the head-neck system at low frequencies (Fig. 7B). It is within this range that active, sensory-driven stabilization is required, which we discuss further in the next section.

Some of our observations highlight that there may be consequences of a flexible head-neck joint and inertial stabilization which are not obviously beneficial. At times, the head became stabilized at an offset from the default level orientation (Fig. 6D), with the constant error of the head angle going uncorrected over multiple stimulus cycles. A similar uncorrected head angle error was reported in a previous study, apparently as result of over-shoot from a step rotation ^46^. We suggest that the overshoot itself may have been caused by the freely spinning head-neck joint. Even without sensory input and stabilizing reflexes, these events would not be expected to occur in other species, where elasticity in the neck motor system likely provides a passive restoring force to correct for static offsets during flight ^59^.

The second requirement for the hoverfly head-neck joint is an ability to switch between the aforementioned passive, free-spinning mode and a mode in which the muscles of the neck motor system exert control over the movement of the head. Active head movements are made during flight, not just around the roll axis, but also around pitch and yaw ^5,47^. Grooming, feeding and other behaviors also require fine motor control of the head. A mechanism should therefore exist to temporarily disengage the neck motor system. Its point of action could be the physiology of the muscles or their mechanical coupling of the head and thorax—a feature which could be resolved with fast in vivo imaging ^60^.

Surprisingly, both of these requirements appear to be met by properties of the head-neck joint in another flying—and hovering—group of insects: the dragonflies and damselflies. The ‘head-arrester’ system found in the adults of all known species of Odonata is an arrangement of muscles and skeletal structures in the neck joint which mechanically lock the head to the thorax ^61,62^. Movement of the head can be selectively enabled by release from the arrested state. The head pivots at a single-point and folds in the connective membranes of the arrester system impart a high degree of flexibility to the joint ^63^. The main purpose of the head-arrester system is thought to be reinforcement of the neck, which is generally very thin compared to the size of the head and a mechanical weak-point ^62,64^. During certain behaviors, such as feeding or tandem flights, the head is arrested in order to prevent injury to the neck ^61,65^.

For agile flight maneuvers, such as chasing, the dragonfly head appears to be free to move and, just as in the hoverfly, inertia acts to stabilize it in a default orientation ^61^. A passive gaze stabilization system may be advantageous in dragonflies and damselflies, since they lack the specialized fast mechanosensory input provided by the halteres in Diptera. The head is also typically larger and of greater mass in dragonflies than in hoverflies, which may help to passively maintain a default orientation of the head even without dynamic movement ^61^. Intriguingly, in the un-arrested state certain contact points between structures in the head-neck joint become physically separated, causing fields of mechanosensory sensilla on their surfaces to be disabled ^62^. These sensilla usually monitor the position of the head relative to the thorax and appear to be involved in flight reflexes and gaze stabilization ^61,62^. Without this proprioceptive information, offsets in the roll angle of the head can go uncorrected during inertial stabilization in dragonflies, just as we and others ^46^ have observed in hoverflies.

The anatomy of the neck-motor system is well-described in dragonflies and blowflies, and they exhibit many fundamental differences to each other ^5,61^—unsurprising, given their evolutionary divergence ^2^. Similar descriptions are unfortunately lacking in hoverflies, and we can only speculate as to how inertial stabilization of the hoverfly head may be selectively enabled and disabled. However work is now underway to provide a detailed anatomical study and to search for a mechanism which may be functionally equivalent to the odonate head-arrester system.

### A hybrid gaze stabilization system with active and passive components

Hoverflies show a remarkably improved gaze stabilization performance at high stimulation frequencies, presumably enabled by a passive, inertial mechanism. An inertia-driven system appears only to operate under high rotational accelerations in hoverflies. At stimulation frequencies below 15 Hz, we observed a gaze stabilization reflex which largely resembles those found in the blowfly and horsefly, whereby sensory input is required. In this lower dynamic range, the halteres play a significant role by sending a forward signal to initiate fast compensatory head movements with low response latency. This reduces the motion of the head—and thus the retinal slip speed—sufficiently to allow the motion vision pathway to also provide feedback signals to the stabilization reflex ^35,66^.

All three families share this general principle of sensory-driven, active stabilization, while hoverflies also exhibit a family-specific adaptation to cope with a higher dynamic range. Without the response latency incurred by sensory transduction, neural processing, and the actuation of muscles in the neck-motor system, an inertial system provides clear benefits during flight maneuvers with particularly high accelerations, such as hovering or departures from hovering. As with the control of flight, passive stability can counterbalance the loss of fast sensory input ^67^. And similar to damselflies and dragonflies, the hybrid system that hoverflies have developed is a prime example of morphological computation ^68,69^ where functional anatomical structures enable the highly effective performance of specific sensorimotor control tasks. The design of energy-efficient, artificial image stabilization systems may take inspiration from this novel biological approach ^70^.

## ACKNOWLEDGMENTS

We thank Dexter Gajjar-Reid for assistance with experiments, Gregor Belušič for providing horseflies, and Kevin Fancourt of Johns Lane Farm for access to land for the collection of hoverflies. This research was supported by an EPSRC Industrial CASE PhD studentship sponsored by the Defense Science and Technology Laboratory (DSTL) to BJH, and an AFRL/AFOSR grant FA9550-14-1-0068 to HGK.

## DATA AVAILABILITY

The data and analysis code generated during this study are available at the Open Science Framework: https://osf.io/bhytv

## AUTHOR CONTRIBUTIONS

Ordered according to main list of authors:

**Conceptualization:** BJH, HGK

**Data curation, validation:** BJH

**Formal analysis:** BJH, FJHH, DAS

**Funding acquisition, resources, administration:** HGK

**Investigation:** BJH, KB

**Methodology:** BJH, KDL, DAS

**Software:** BJH, FJHH, KDL, DAS

**Supervision:** KDL, HGK

**Visualization:** BJH, FJHH

**Writing – original draft:** BJH, HGK

**Writing – review & editing:** BJH, KB, FJHH, KDL, DAS, HGK

## MATERIALS AND METHODS

### Animal collection and preparation

Wild-type, adult female flies of indeterminate age were used for all experiments. Blowflies, *Calliphora vicina*, were collected from a colony raised in lab conditions at 20°C, on a 12:12 hour dark:light cycle. Wild horseflies, *Tabanus bromius*, were caught in fields in Buckinghamshire, UK and near Ljubljana, Slovenia. Wild hoverflies, *Episyrphus balteatus* and *Eristalis tenax*, were caught in Buckinghamshire, UK. Hoverflies raised in commercial colonies were also used, transported as pupae: *Eristalinus aeneus* from Bioflytech SL, Spain, and *Episyrphus balteatus* from Katz Biotech AG, Germany. Prior to experiments, animals were kept in net cages with conspecifics. Individual flies were collected from their cage and cooled on ice in a vial. A cardboard tether was attached to the pro-thorax using beeswax. The tether was oriented to give an approximately 0°attitude of the body during tethered-flight. For experiments with the halteres removed, the shaft of the halteres was severed as close as possible to its base using sharp micro-dissection scissors. Normal wing-stroke, leg-tuck and head movements were verified before experiments. Although we considered testing anesthetized or sacrificed animals, finding a lack of inertial stabilization in this condition could have a number of possible causes, such as a disabled mechanism for switching to a passive head-neck joint.

### Experimental setup

Tethered animals were secured to a step-motor which was controlled by a micro-stepping driver (P808, Astrosyn). The motor step resolution used was either 5000 or 3200 steps per revolution, for 0–10 Hz or 15–25 Hz oscillations, respectively. The motor driver was controlled through Matlab (R2014a, Mathworks) via a DAQ (NI-6025E, National Instruments). A hemispherical false horizon made of black-painted plastic, approximately 50 mm diameter, was positioned beneath the animal with the top edge close to the eye equator. A slightly larger diameter translucent white plastic hemisphere was positioned above the fly to form a light diffuser which encompassed the horizon (Fig. 1A). Illumination was provided by four light guides (KL 1500, Schott). Luminance at the position of the animal was measured to be 500 Cd m^−2^. A small opening in the front of the horizon permitted a head-on view of the animal. Airflow was applied continuously during experiments to encourage flight.

Two high-speed cameras were used to record experiments: one for shorter experiments (Fastcam SA3, Photron) with a 100 mm macro lens (Zeiss), and one with higher storage capacity for longer experiments (Phantom v211, Vision Research) with a 180 mm macro lens (Sigma). Aperture sizes were adjusted between *f* /3.5–5.6 depending on the length of the animal and depth-of-field required. Frame-rates up to 1200 fps were chosen according to the length of the experiment and the stimulus frequency, ensuring at least 1 frame per 2° of rotation.

### Stimulus protocol

The chirp stimulus time-series was defined as:

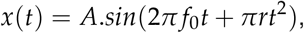

where *A* is the oscillation amplitude (30°), *f*_0_ is the initial frequency (0 Hz), *t* is the time vector, and *r* is the chirp rate—the rate of change in frequency—over the time interval, *T* (10 s):

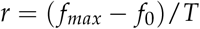

A positive and a negative chirp rate were used within each experiment:

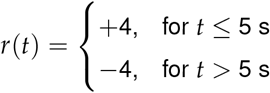

with a maximum frequency, *f*_*max*_, of 20 Hz. Experiments using constant-frequency stimuli varied in length and number of cycles, from 3 cycles at 0.06 Hz to 250 cycles at 25 Hz. Experiments using 15–25 Hz stimuli required an initial ramp in amplitude to overcome the inertia of the step motor: the amplitude reached ±30° within 2 s, and 10 s of subsequent cycles were analyzed per experiment.

### Video analysis

Recorded experiments were analyzed automatically to extract the roll angles of the head and the cardboard tether in each video frame. Analysis was carried out in Labview (v2013, National Instruments) using a modified version of a previously-developed custom template-matching method ^71^. Only experiments in which the animal flew continuously for all stimulus cycles were analyzed. Subsequent analysis of roll angle time-series was carried out in Matlab (2020b, Mathworks).

### Maximum stimulus velocity

For constant-frequency sinusoidal oscillations, the angular velocity of the stimulus varied throughout each cycle. For plots of slip-speed distribution (Fig. 4, Fig. S1) we marked the theoretical maximum slip-speed experienced with no stabilization effort (i.e. head angle = thorax angle), which we calculated as the maximum angular velocity of the stimulus in each cycle:

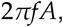

where *f* is the oscillation frequency and *A* is the oscillation amplitude.

### Motion blur limit

The retinal slip speed at which motion blur occurs was approximated from a rule-of-thumb of one photoreceptor acceptance angle per response time ^72^. With an estimated range of acceptance angles of 1–2° for the species studied ^73,74^ and a response time of 10 ms, motion blur would be expected to begin to degrade visual information at slip speeds around 100–200°s^−1^ and higher. Note that this does not imply an upper limit to useful motion vision—responses in motion-sensitive neurons in Diptera have been recorded at greater image velocities ^17^.

### Head-neck model

A previously-developed model of the dynamics of blowfly gaze stabilization ^75^ was modified to include only the passive physical properties of the head and neck. The following equation of motion for the head was solved at discrete time intervals:

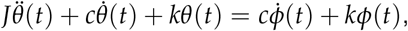

where *θ* is the roll angle of the head, *ϕ* is the roll angle of the thorax (determined by the chirp stimulus time-series described above), *k* and *c* are the torsional spring and damping constants of the head-neck joint, respectively, and *J* is the moment of inertia of the head, defined for a thin-walled spherical shell (approximating the hoverfly head) as:

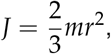

where *m* is the mass of the sphere and *r* is its radius.

The following values for physical parameters were used: *m* = 10 × 10^−6^ kg, *r* = 0.002 m, *J* = 2.66 × 10^−11^ kg m^2^, *k* = 1 × 10^−8^ N m deg^−1^, *c* = 1 × 10^−9^ N m s deg^−1^. The values chosen for *k* and *c* were one order of magnitude smaller than those estimated for the blowfly ^75^, in order to investigate the proposed low stiffness and damping of the hoverfly head-neck joint.

## SUPPLEMENTARY INFORMATION

**Figure S1.**
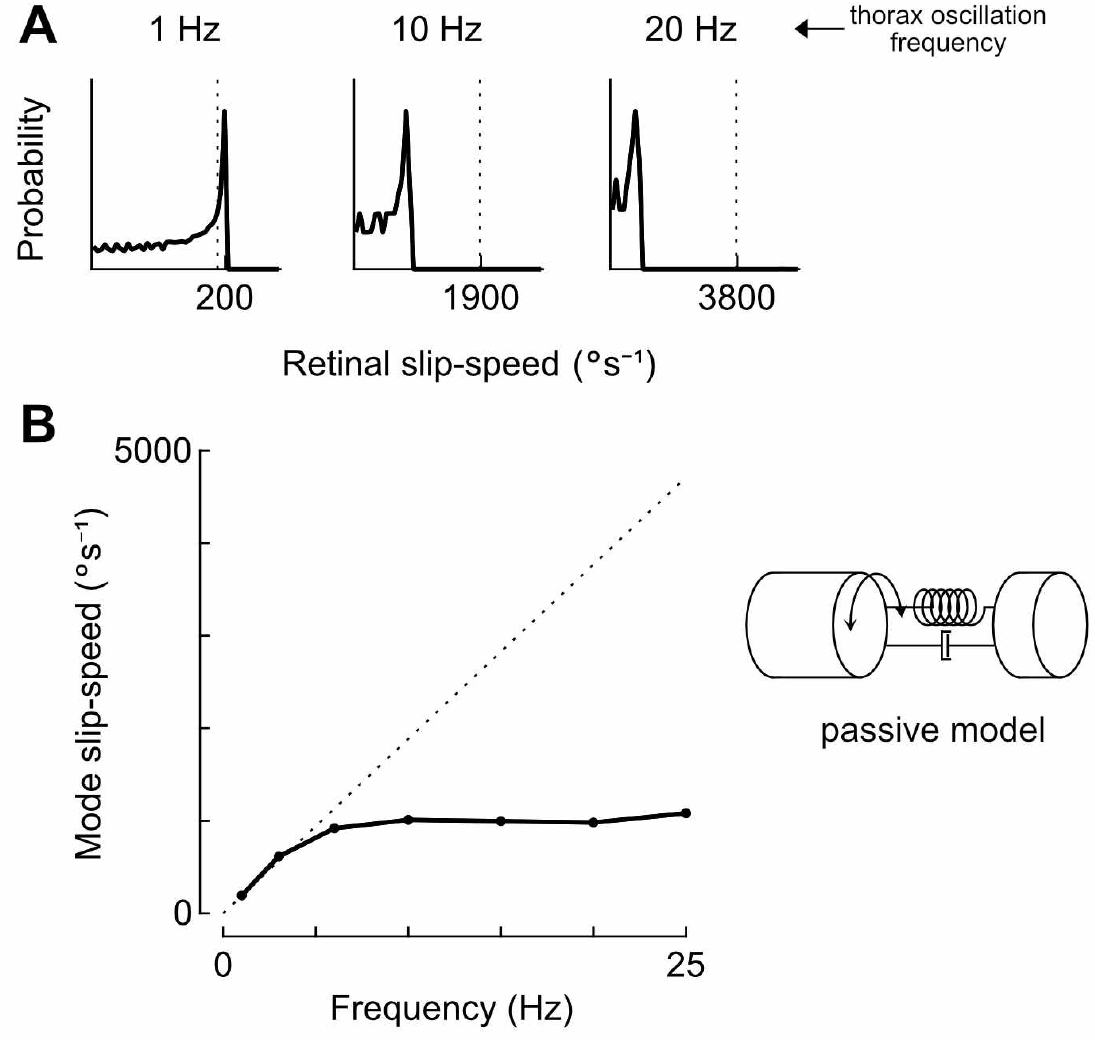
Slip-speed distribution at different frequencies for the head-neck model. **A:** Normalized probability distribution of visual slip experienced by the passive model head shown in Fig. 7, during simulated constant-frequency sinusoidal oscillations. Vertical dashed line indicates theoretical maximum slip-speed experienced with no stabilization effort (i.e. head angle = thorax angle). **B:** Mode (peak) values of the probability distributions of visual slip experienced by the passive model head during simulated constant-frequency sinusoidal oscillations.

**Movie 1. High-speed video of *C. vicina* chirp experiment** https://osf.io/qyc3m

**Movie 2. High-speed video of *T. bromius* chirp experiment** https://osf.io/sntdf

**Movie 3. High-speed video of *E. aeneus* chirp experiment** https://osf.io/d3njt

**Movie 4. High-speed video of *E. balteatus* chirp experiment** https://osf.io/4zrpa

**Movie 5. High-speed video of *E. tenax* chirp experiment** https://osf.io/s6kj3

